# DNA damage leads to microtubule stabilisation through an increase in Golgi-derived microtubules

**DOI:** 10.1101/2022.08.29.505705

**Authors:** Aishwarya Venkataravi, Mayurika Lahiri

## Abstract

The site of nucleation strongly determines microtubule organisation and dynamics. The centrosome is a primary site for microtubule nucleation and organisation in most animal cells. In recent years, the Golgi apparatus has emerged as a site of microtubule nucleation and stabilisation. The microtubules originating from Golgi are essential for maintaining Golgi integrity post-Golgi trafficking, establishing cell polarity and enabling cell motility. Although the mechanism of nucleation and functional relevance of the Golgi-nucleated microtubule is well established, its regulation needs to be better studied. In this study, we report that DNA damage leads to aberrant Golgi structure and function accompanied by reorganisation of the microtubule network. Characterisation of microtubule dynamics post DNA damage showed the presence of a stable pool of microtubules resistant to depolymerisation by nocodazole and enriched in acetylated tubulin. Investigation of the functional association between Golgi dispersal and microtubule stability revealed that the Golgi elements were distributed along the acetylated microtubules. Microtubule regrowth assays showed an increase in Golgi-derived microtubule post DNA damage. Interestingly, reversal of Golgi dispersal reduces microtubule stabilisation. Altered intracellular trafficking resulting in mislocalisation of cell-cell junction proteins was observed post DNA damage. We propose that the increase in stable microtubules deregulates intracellular trafficking, resulting in cell polarity changes. This study would thus be the first to demonstrate the link between Golgi dispersal and microtubule reorganisation orchestrating changes in cell polarity.

## Introduction

Microtubules (MTs) are one of three dynamic polymers that constitute the cell’s cytoskeleton. Microtubules are formed by end-to-end polymerisation of α/β- tubulin heterodimers which come together to form a hollow cylindrical structure. They are polarised filaments with the minus (-) end anchored at the nucleating centre and the plus (+) end radiating towards the cell membrane. The centrosome is the primary microtubule organising centre in mammalian cells, followed by the Golgi apparatus. The organisation and dynamics of the MT arrays are determined by their site of nucleation. Golgi-derived microtubules (GDMTs) are dynamically different from centrosomal microtubules. GDMTs are more stable and enriched in tubulin acetylation and detyrosination (Chabin-Brion et al., 2001; Skoufias et al., 1990; Thyberg and Moskalewski, 1993). They contribute to cell asymmetry by facilitating polarised trafficking toward the cell’s leading edge during cell migration (Vinogradova et al., 2009).

Microtubules play myriad functions inside the cell, and microtubule dynamics are essential in regulating these functions. A complex signalling mechanism is in place to translate externals cues to a cellular response through changes in microtubule dynamics. A growing body of literature emphasises the role of microtubules in the cellular response to stress such as hypoxia, metabolic stress, mechanical stress and genotoxic stress (Parker et al., 2014).

A decrease in microtubule polymerisation is observed in anoxic conditions, while increased polymerisation is observed under hypoxia(Hu et al., 2010). The response to hypoxia has been reported to occur through GSK3β, resulting in Rab11-dependent trafficking of α6β4-integrin(Yoon et al., 2005). Under anoxic conditions, MAP4 phosphorylation leads to microtubule stabilisation (Hu et al., 2010). The dynein light chain has been shown to regulate mitochondrial permeability through its interaction with voltage-gated anion channels(Fang et al., 2011). Microtubules are known to respond to metabolic stress and have been implicated in regulating cellular metabolism (Bershadsky and Gelfand, 1981). AMPK, a metabolic stress sensor, phosphorylates CLIP170 and affects microtubule dynamics(Nakano et al., 2010). In neuronal cells, axonal microtubule growth is inhibited upon metabolic stress in an AMPK-dependent manner (Williams et al., 2011). Nutrient starvation has also been shown to lead to microtubule hyperacetylation and has been associated with starvation-induced autophagy (Geeraert et al., 2010).

DNA damage response (DDR) is a well-orchestrated signalling network which, in response to DNA damage, leads to the detection of DNA lesions, cell cycle arrest, transcriptional activation of DNA repair genes and, ultimately, DNA repair (Jackson and Bartek, 2009). Classical studies contributing to our understanding of DDR majorly involve nuclear processes. The cytoplasmic effect of DDR, especially concerning the cytoskeleton, is recently being explored.

One of the initial studies linking DDR to cytoskeleton showed that p53, an essential DDR protein, is transported to the nucleus via microtubules in response to DNA damage (Giannakakou et al., 2000). Later a study in 2014 reported that upon DNA damage, DNA-PK phosphorylates GOLPH3 leading to Golgi dispersal through actin modulation (Farber-Katz et al., 2014). GOLPH3 is a Golgi membrane protein that interacts with the unconventional MYO18A-F-actin complex to maintain the perinuclear ribbon morphology of the Golgi apparatus (Dippold et al., 2009). Phosphorylation by DNA-PK leads to increased interaction between GOLPH3-MYO18A which adds a tensile force on the Golgi, thereby dispersing it throughout the cytoplasm (Farber-Katz et al., 2014). The following year, Tito Fijo’s group showed that microtubule poisons interfere with the transport of DNA repair proteins to the nucleus, delaying DNA repair (Poruchynsky et al., 2015). A more recent study by Mi Ryu reports microtubule stabilisation upon induction of genotoxic stress. An increase in tubulin acetylation by αTAT1(α-Tubulin acetyltransferase) is essential for activating the S and G2/M checkpoints post-DNA damage (Ryu and Kim, 2020). Although this was the first study to report a direct association of microtubule dynamics with DNA damage, it did not delve into how DNA damage might lead to microtubule stabilisation. A more recent study reported that DNA damage leads to a transient change in microtubule dynamics which is necessary for DNA repair through the non-homologous end joining (NHEJ) pathway. They observed an increase in the polymerisation of centrosomal microtubules through the activation of DNA-PK but did not comment on the stability of the microtubules (Ma et al., 2021).

In this study, we report that DNA damage leads to dramatic reorganisation of the microtubules and their dynamics through increased GDMTs. Treatment with DNA damaging agents led to microtubule stabilisation accompanied by dispersal of Golgi, as previously reported. Further investigation revealed that the changes in Golgi morphology were essential for microtubule stabilisation. DNA damage-induced Golgi dispersal led to an increase in GDMTs, which are stable. Interestingly, an increase in genomic instability induced stable microtubules and Golgi-derived microtubules were observed in MCF10CA1a cells in comparison to MCF10A. We also report impaired intracellular trafficking leading to a mislocalisation of cell-cell junction proteins, possibly due to altered microtubule dynamics.

## Results

### DNA damage leads to microtubule stabilisation

Non-tumorigenic MCF10A cells were treated with a sub-lethal dose of 2mM NEU (Bodakuntla et al., 2014) or 10J/m^2^ UV(Fong et al., 2010) to induce DNA damage and were analysed for microtubule stabilisation. Microtubule stabilisation was tested using two complementary methods – one, checking for resistance to depolymerisation induced by nocodazole treatment (Xu et al., 2017), and the other was monitoring the +TIP dynamics using the EB3 construct (Miller et al., 2009). MCF10A cells with and without DNA damage were treated with nocodazole (2.5μg/ml, 20mins) and stained for α-tubulin to assess the extent of microtubule depolymerisation. We observed that cells post DNA damage showed resistance to nocodazole treatment **(Fig 1a-b)**. Nocodazole treatment revealed a subset of microtubules which were hyper stable and resistant to depolymerisation, indicating a change in microtubule dynamics.

**Figure 1:**
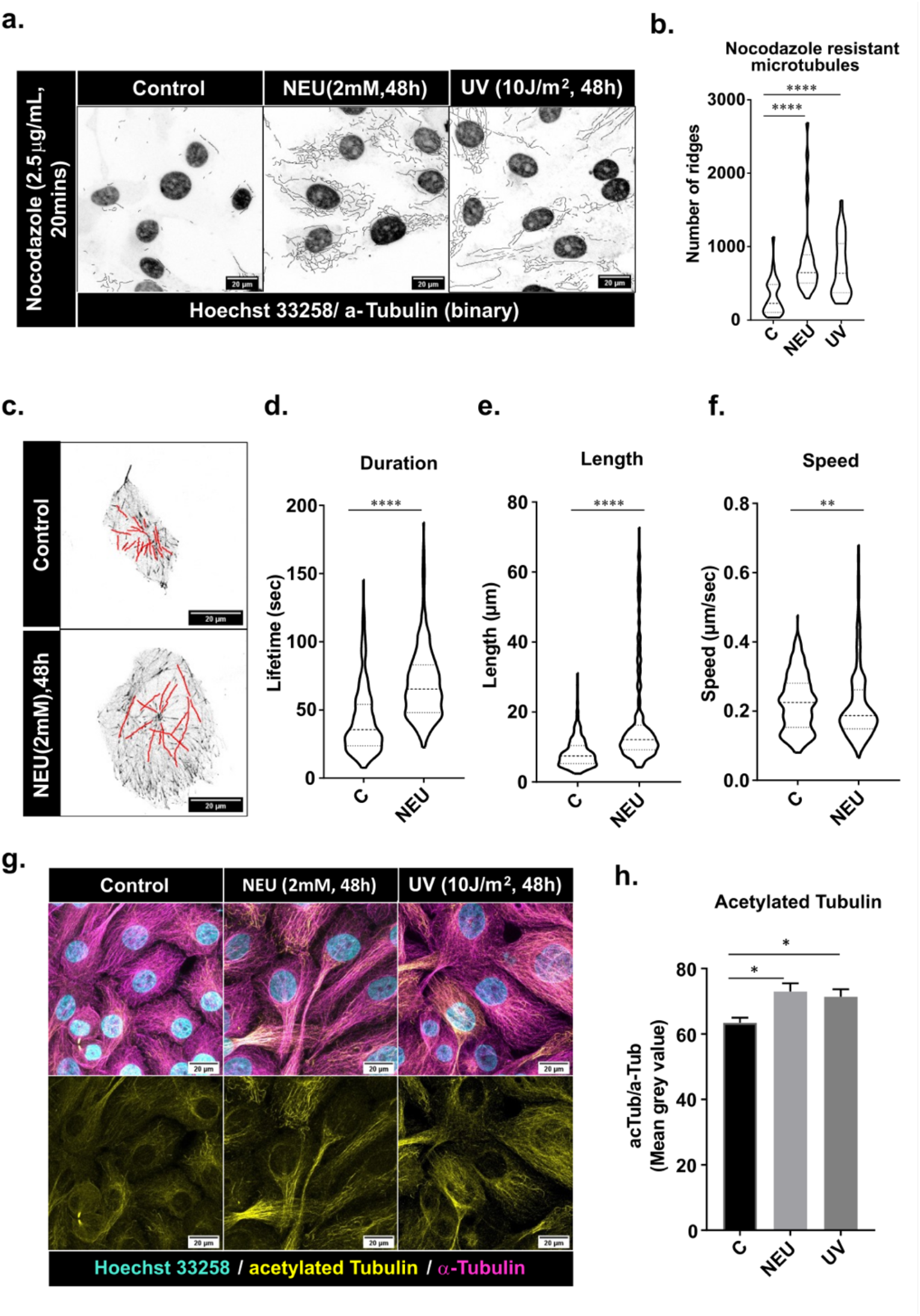
MCF10A cells showed an increase in microtubule stability post-DNA damage. (**a**) NEU (2 mM, 48 hrs) or UV (10 J/m^2^) damaged cells, when treated with nocodazole, revealed a subset of microtubules resistant to depolymerisation that were represented as binary images. (**b**) The nocodazole-resistant MTs were quantified using ImageJ ridge detection tool (N=3, n=140 cells). Asterisks indicate Mann-Whitney U test significance values; **** p < 0.0001. (**c**) EB3 assay was performed on cells treated with and without 2 mM NEU for 48 hrs, and live-cell imaging was performed. A few EB3 comets are shown. The EB3 comets were manually tracked and quantified using MtrackJ plugin on ImageJ to show comet (**d**) duration, (**e**) length and (**f**) speed (N=3, n=120 cells, 25 MT tracks per cell). (**g**) MCF10A cells treated with NEU (2mM, 48h) and UV (10J/m^2^) were stained for α-tubulin (magenta), acetylated tubulin (yellow), and Hoechst 33258 (cyan), showed increased tubulin acetylation post DNA damage. (**h**) Bar graph representing the quantification of acetylated tubulin (N=3, n=125 cells). Statistics analysis was performed for all the experiments using the Mann-Whitney U test with significance values **** p < 0.0001; ** p< 0.01; * p < 0.05.

Further, we used EB3 dynamics as a surrogate for assessing microtubule growth dynamics. Cells with and without DNA damage were transfected with the EB3 construct, and the EB3 comets were imaged. NEU-treated cells appeared to have longer microtubule tracks than untreated cells **(Fig 1c).** Quantifying EB3 dynamics showed a significant increase in MT lifetime and length. At the same time, a decrease in speed was observed in the NEU-treated cells compared to the untreated control, indicating MT stabilisation **(Fig 1d-f)**. Tubulin acetylation is a tubulin PTM which is often associated with stable microtubules. Hence, cells were stained with α-tubulin to check if DNA damage led to changes in tubulin acetylation. A significant increase in K40 tubulin acetylation was observed in both NEU and UV-treated MCF10A cells compared to the untreated control cells **(Fig 1 g** and **h)**. We also observed that post DNA damage, the microtubules were arranged in parallel arrays, compared to the radial arrays observed in the untreated cells **(Supp Fig 1 a-c)**. In addition, a dramatic change in the morphology of the cells was observed after treatment with NEU. The cells appeared to be more elongated and larger, where the NEU-treated cells showed significant increases in the cell area, nucleo-cytoplasmic ratio, circularity and aspect ratio **(Supp Fig 1 d-g),** which might be attributed to the changes in the microtubule network. This phenotype was further confirmed in the HEK293 cell line. The NEU dose was first standardised for the HEK293 cells where the cells were treated with 0.5 mM, 1 mM and 2 mM NEU and stained for phosphorylated DNA-PK (T2609) to obtain a sublethal dose of NEU with sufficient DNA damage **(Supp Fig 2 a-b)**. NEU dose of 0.5 mM dose was used for further experiments. An increase in nocodazole-resistant microtubules, as well as an increase in tubulin acetylation, were observed when HEK293 cells were treated with 0.5 mM NEU for 48 hrs and 10 J/m^2^ UV for 48 hrs **(Supp Fig 2 c-f)** thus demonstrating that the changes in microtubule dynamics are not cell line specific.

### Activation of DNA-PK precedes Golgi dispersal, followed by the appearance of a stable pool of microtubules

To decipher how DNA damage might lead to changes in microtubule dynamics, we investigated the activation of DNA-PK and Golgi dispersal following DNA damage. Since Golgi has a close association with microtubules, we proposed that changes in microtubule dynamics might be through Golgi dispersal. Activation of DNA-PK (T2609) and dispersal of Golgi were detected 48 hrs post-NEU and UV treatment **(Supp Fig 3 a-h),** thus implying that Golgi dispersal post-NEU and UV treatment might be through the DNA-PK-GOLPH3 pathway (Farber-Katz et al., 2014). The presence of active DNA-PK foci at 48 hrs demonstrated that DNA damage lesions have not been repaired and that the change in microtubule dynamics may be in response to the DNA damage. To investigate the temporal dynamics of DNA-PK activation, Golgi morphology changes and tubulin acetylation occurring post DNA damage, cells were treated with 2 mM NEU for different time durations and analysed. Phosphorylation of DNA-PK (T2609) was observed as early as 10 mins, while Golgi dispersal was observed from 4 hours post-NEU treatment **(Fig 2 a-b; Supp Fig 4 a-b)**. Both these events preceded the presence of nocodazole-resistant microtubules and tubulin acetylation, that were observed 18 hrs post-NEU treatment **(Fig 2 c-f)**.

**Figure 2:**
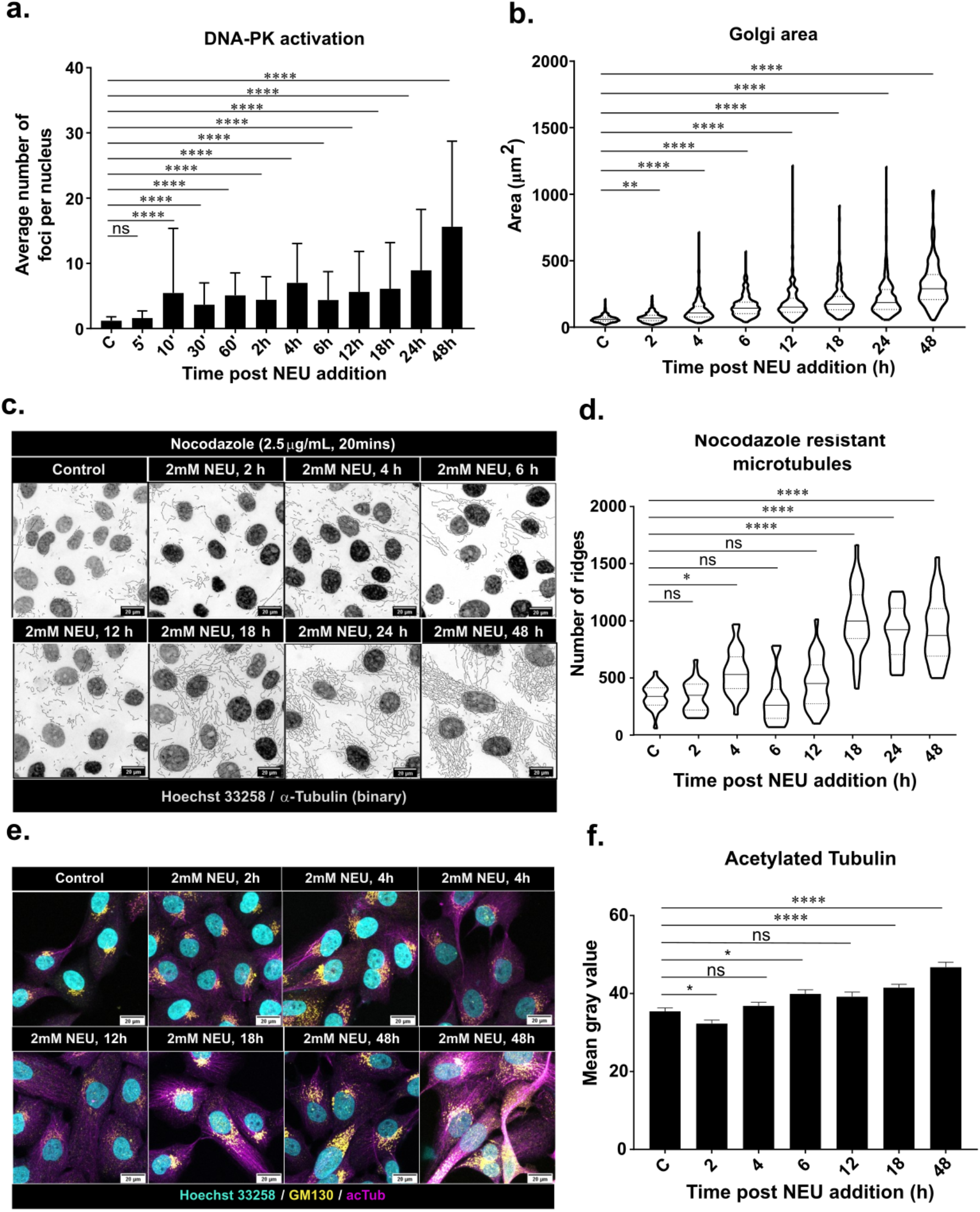
The activation of DNA-PK and Golgi dispersal precedes microtubule stabilisation. MCF10A cells were treated with 2 mM NEU and checked for (**a**) activation of DNA-PK at threonine 2609 at different time points. A significant increase in pDNA-PK foci was observed from 10 mins post NEU treatment which remained active till 48 hrs (N=3, n=420 cells). (**b**) Golgi dispersal was observed 4 hrs post DNA damage (N=3, n=220). (**c**) the appearance of nocodazole-resistant microtubules at 18hrs after NEU treatment was depicted as binary images of tubulin and nucleus (both in grey) that were quantified in (**d**) using a ridge detection tool (N=3, n>10 fields per timepoint). (**e**). An increase in tubulin acetylation (magenta) was observed at 18 hrs in cells with dispersed Golgi (yellow). Hoechst 33258 (cyan) was used to stain the nucleus in the cells (N=3, n=290 cells). (**d**) Bar graphs showing the quantification of acetylated tubulin in the cells. Kruskal-Wallis test for significance was performed on all the data **** p < 0.0001, **p< 0.01, *p< 0.05 ns – nonsignificant.

To test whether activation of DNA-PK is required for microtubule stabilisation, cells were treated with DMNB, a small molecule inhibitor against the catalytic subunit of DNA-PK (Durant and Karran, 2003). A significant reduction in tubulin acetylation (**Fig 3 a-b**) and nocodazole-resistant microtubules (**Fig 3 c-d**) were observed in cells treated with DMNB following 2mM NEU treatment for 48 hrs compared to the untreated control cells. Inhibition of DNA-PK was previously reported to revert DNA damage-induced Golgi dispersal (Anandi et al., 2017; Durant and Karran, 2003; Farber-Katz et al., 2014). DNA damage-induced Golgi dispersal has been shown to involve the actin cytoskeleton, and its depolymerisation to revert the dispersal of Golgi (Dippold et al., 2009; Farber-Katz et al., 2014). To assess whether activation of DNA-PK leads to microtubule reorganisation by altering Golgi distribution, Latrunculin A (LatA, an actin depolymerising agent) was used to condense Golgi in the presence of DNA damage. LatA treatment led to a significant reduction in acetylated tubulin (**Fig 3 e-f)** and nocodazole-resistant microtubules (**Fig 3 g-h)**, similar to the inhibition of DNA-PK. Thus, it may be inferred that the stabilisation of microtubule post DNA damage is mediated through the DNA-PK-Golgi axis.

**Figure 3:**
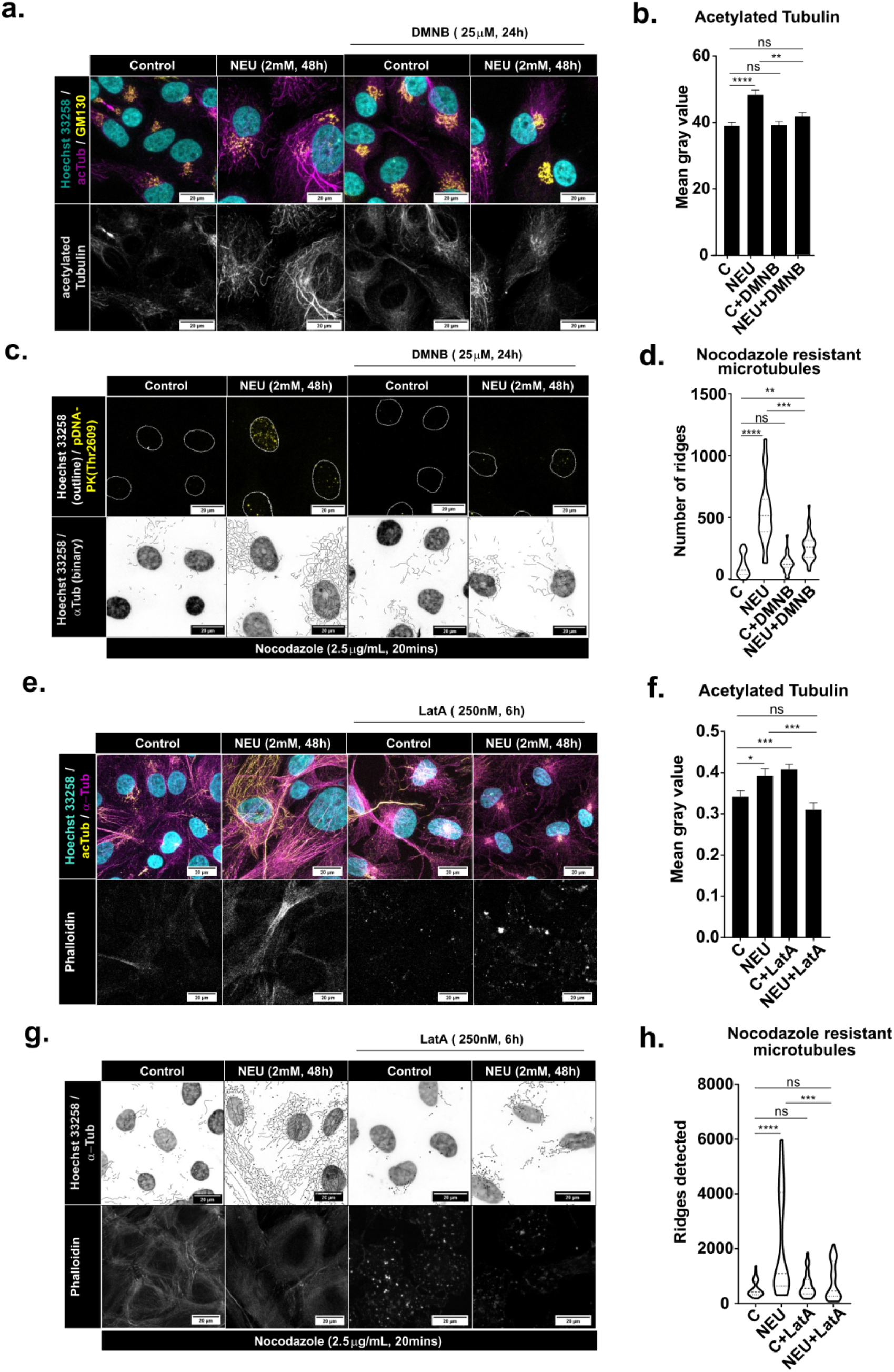
DNA damage-induced microtubule stabilisation is through activation of DNA-PK and Golgi dispersal. Cells were treated with 2 mM NEU for 48 hrs and to one set DMNB (25μM, two doses 12 hrs each) was added to inhibit DNA-PK. (**a**) The cells were stained for GM130 (yellow) that marked the Golgi, Hoechst 33258 (cyan) for the nucleus and acetylated tubulin (magenta). Inhibition of DNA-PK using DMNB post-NEU treatment showed a reduction in the acetylated tubulin in DMNB and NEU-treated cells compared to cells with only NEU treatment (N=3, n=100 cells). (**b**) The intensity of acetylated tubulin was quantified and represented as bar graphs. (**c**) 2.5 μg/ml nocodazole was added to 2 mM NEU-treated cells for 48 hrs with or without DMNB and checked for pDNA-PK (Thr 2609) foci (yellow) and α-tubulin (binary image) (N=3, n>10 fields per treatment). (**d**) violin plots representing nocodazole-resistant microtubules that were quantified using a ridge detection tool. (**e**) Latrunculin A (Lat A) was added at 250 nM for 6 hrs to the NEU-treated (2mM, 48 hrs) cells and control cells and stained for Hoechst 33258 (cyan) for the nucleus, acetylated tubulin (yellow) and α-tubulin (magenta). Phalloidin was used as a positive control for Lat A treatment (N=3, n=110 cells). Depolymerisation of F-actin by Lat A led to a reduction in tubulin acetylation that was quantified in (**f**) and represented as bar graphs. (**g**) Lat A (250 nM, 6 hrs) was added to nocodazole (2.5 μg/ml, 20 mins) treated control and NEU (2 mM, 48 hrs) treated cells and stained for Hoechst 33258 and α-tubulin. Phalloidin was used as a positive control for Lat a treatment (N=3, n>10 fields per treatment). (**h**) Violin plots representing nocodazole-resistant MTs that were quantified using a ridge detection tool. Statistical analysis was performed using Kruskal-Wallis test with significance values; **** p < 0.0001, ***p< 0.001, **p< 0.01 and ns – non-significant.

### DNA damage leads to an increase in Golgi-derived microtubules

Golgi and microtubules are functionally dependent on each other. Microtubules play an essential role in maintaining Golgi structure (Vinogradova et al., 2012). Treating cells with microtubule poisons displays a fragmented and dispersed Golgi (Thyberg and Moskalewski, 1993; Wehland et al., 1983). On the other hand, a significant proportion of microtubules in mammalian cells originate from Golgi, and there is increasing evidence that these microtubules are stable. We hypothesised that the stable pool of microtubules which appear post DNA damage are non-centrosomal microtubules that nucleated at the Golgi apparatus. In the cells treated with NEU, we observed that the stable acetylated microtubules were associated with the dispersed Golgi apparatus, which indicated that they may have nucleated from the Golgi **(Fig. 4a)**. To confirm whether there is an increase in GDMTs, microtubule regrowth assay was performed following treatment of cells with 2.5 μg/ml nocodazole. A time-course was performed to check for complete depolymerisation of the microtubules. A treatment of 2.5 μg/ml nocodazole for 3 hrs resulted in almost complete depolymerisation of microtubules which was used for the washout assay **(Fig. 4b)**. Close observation of cells recovering from microtubule depolymerisation showed that a significant number of microtubules were nucleated from the Golgi post-NEU treatment **(Fig.4c-d)**.

**Figure 4:**
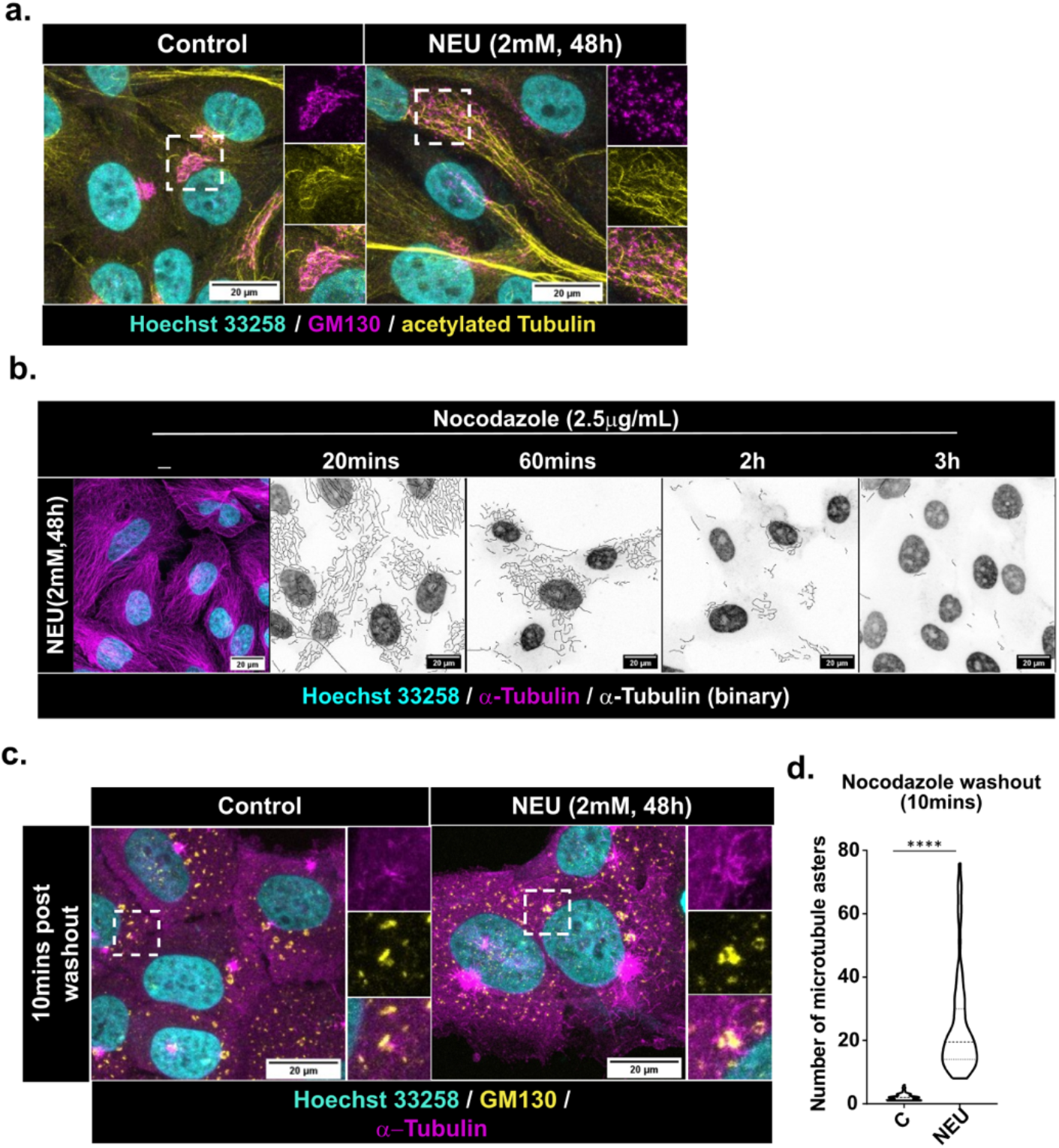
NEU-induced DNA damage leads to an increase in Golgi-derived microtubules. (**a**) Untreated and 2 mM NEU treated (48 hrs) MCF10A cells were stained for acetylated tubulin (yellow), GM130 (magenta) and Hoechst 33258 (cyan) to check for Golgi elements distributed along the acetylated microtubules (N=2, n=75 cells). (**b**) Representative images of NEU-treated cells incubated with 2.5 μg/mL nocodazole for 20 mins, 60 mins, 2 hrs and 3 hrs are shown. Nocodazole dose of 2.5μg/mL for 3 hrs led to complete depolymerisation of microtubules in NEU-treated cells. (**c**) A Nocodazole washout assay was performed to determine changes in the nucleation of microtubules from the Golgi. Cells were treated with 2.5μg/mL for 3h, then washed with cold PBS and supplemented with fresh media. Microtubules were allowed to recover at 37°C and fixed and stained for α-tubulin (magenta), GM130 (yellow) and Hoechst 33258 (cyan). An increase in microtubule nucleation at Golgi was observed 10 mins post nocodazole washout, which was quantified in (**d**) by manually counting the number of nucleation sites (N=3, n=70). Asterisks indicate Mann-Whitney U test significance values; **** p < 0.0001.

### Golgi-derived microtubules are associated with genome instability

DNA damage-induced Golgi dispersal has been proposed to be an adaptive mechanism in response to genotoxic insults through altering the trafficking of proteins(Buschman et al., 2015; Farber-Katz et al., 2014). Given the importance of the Golgi-microtubule association in intracellular trafficking, we hypothesised that GDMTs might contribute to the survival of cells with genomic instability. To ascertain the association of an increase in GDMT to genome instability, we compared the levels of GDMTs in a malignant cell line with the non-tumorigenic MCF10A. The MCF10A isogenic cell line series was used to investigate the association for a valid comparison. Anandi et al. have shown that MCF10AT1 and MCF10CA1a have an aberrant Golgi structure due to atypically active DNA-PK, which confirmed that the DNA-PK-Golgi axis is active in the cell line (Anandi et al., 2017). Thus, it was interesting to study if the malignant cell line, MCF10CA1a, had higher levels of stable microtubules and Golgi-derived microtubules compared to MCF10A cells.

It was observed that MCF10CA1a had increased levels of tubulin acetylation **(Supp Fig.5 a and b)**. Additionally, increased levels of nocodazole-resistant microtubules were detected at 2.5mg/ml nocodazole incubated for 20 mins **(Supp Fig.5 c and d)**. For studying microtubule regrowth in MCF10CA1a cells, the nocodazole dose was standardised, and a dose of 2.5mg/ml for 3h was found appropriate **(Supp Fig.5 e)**. MCF10CA1a was observed to have a higher number of microtubules nucleated at the Golgi **(Fig.5 a-b)**. Hence, MCF10CA1a, a malignant cell line with inherent DNA damage, has more stabilised microtubules and Golgi-derived microtubules.

**Figure 5:**
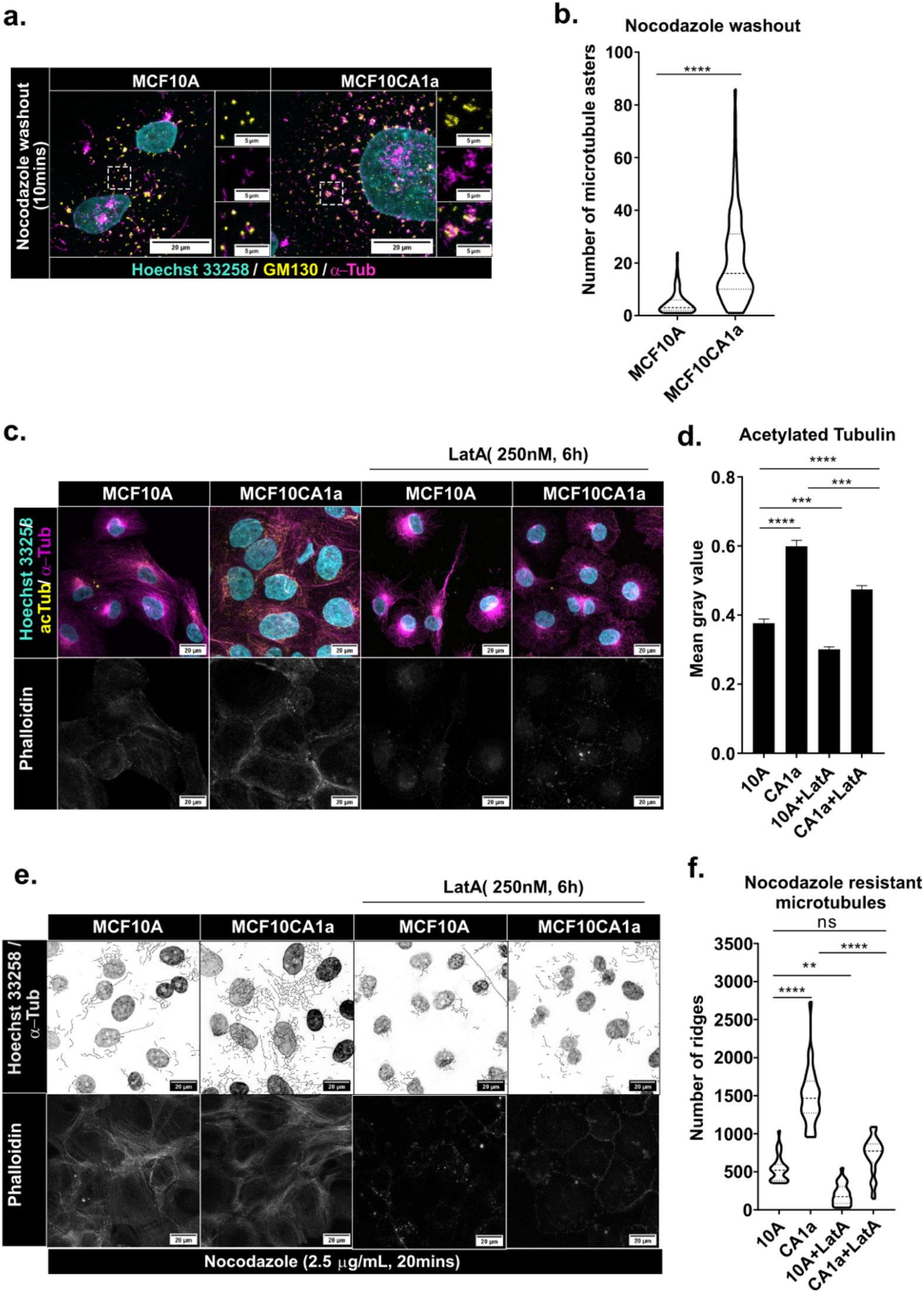
The malignant MCF10CA1a cells have higher levels of Golgi-derived microtubules. MCF10A and MCF10CA1a were stained for microtubules and Golgi post nocodazole washout as previously described. **a.** A higher number of microtubules (magenta) were observed to be nucleated from Golgi (yellow) in MCF10CA1a in comparison to MCF10A, which was quantified by counting the nucleation sites and plotted in (**b.**). Asterisks indicate Mann-Whitney U test significance values; **** p < 0.0001. Microtubule stabilisation in MCF10CA1a was seen to be dependent on Golgi dispersal. Treatment with Lat A (250nM, 6h) led to a reduction in tubulin acetylation (**c. and d.**) and nocodazole-resistant microtubules (**e. and f.**) in MCF10CA1a cells. Asterisks indicate Kruskal-Wallis test significance values; **** p < 0.0001, ***p< 0.001, **p< 0.01 and ns – non-significant.

To assess if the microtubule dynamics in MCF10CA1a are linked to Golgi dispersal, we checked for stable microtubules following Latrunculin A (Lat A) treatment. Upon Lat A treatment, a reduction in tubulin acetylation and nocodazole-resistant microtubules were observed in the MCF10CA1a cells compared to the MCF10A cells **(Fig.5 c-f)**. Therefore, we conclude that microtubule stabilisation in MCF10CA1a cells is also through the DNA-PK-Golgi axis.

### DNA damage results in impaired intracellular trafficking

Golgi and microtubule dynamics are fundamental for transporting cargo from the endoplasmic reticulum (ER) to the destined cellular compartments. Studies have reported that DNA damage-induced Golgi dispersal leads to an impaired intracellular trafficking (Anandi et al., 2017; Farber-Katz et al., 2014). In this study, we wanted to investigate whether NEU treatment leads to impaired intracellular trafficking, thereby affecting cell polarity. RUSH assay with GPI (Glycosylphosphatidylinositol) anchored EGFP as reporters were used for testing for defects in intracellular trafficking post-NEU induced DNA damage (Boncompain et al., 2012). A delayed transport of GPI-EGFP to the plasma membrane was observed, indicating a defect in Golgi trafficking **(Fig. 6a-b).** A dispersed Golgi could also result in defective transport of the cargo to the Golgi. To test this, we used the RUSH construct with mannosidase-II tagged with EGFP (ManII-EGFP) as a cargo. A delayed accumulation of ManII in Golgi was also observed in NEU-treated cells, indicating a defective ER to Golgi transport **(Supp Fig. 6a-b).** To examine if the defects in trafficking affected the localisation of cell polarity proteins, immunostaining of cells with E-cadherin, β-catenin, and α3-integrin showed either mislocalisation of these proteins in the cytoplasm or diffused staining at the junctions **(Fig. 6c-e)** which was quantified **(Fig. 6f-h)** by measuring the CTCF (Corrected Total Cell Fluorescence) of the cytoplasm.

**Figure 6:**
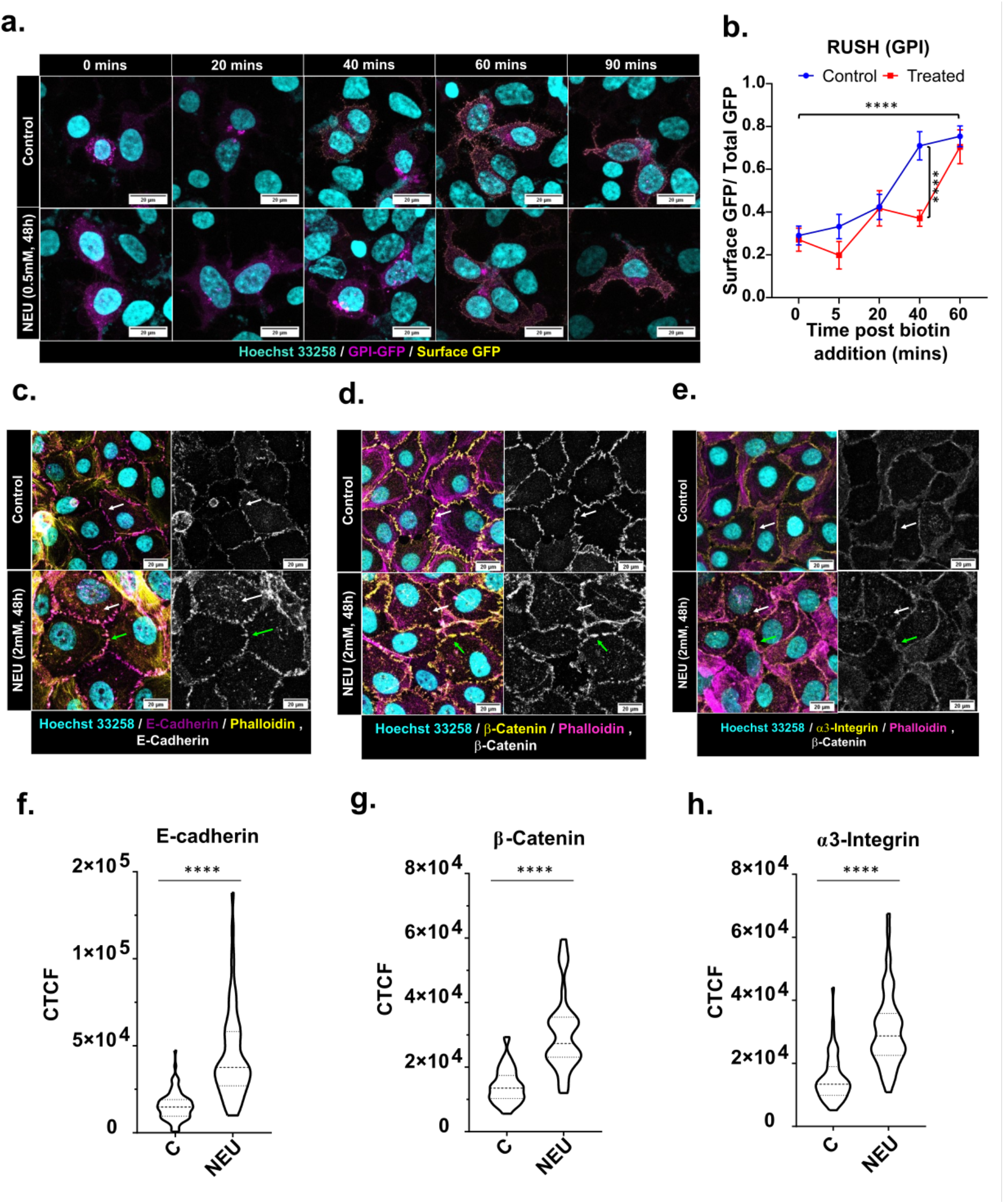
NEU-induced DNA damage led to impaired intracellular trafficking. (**a**) Representative images of cells expressing GPI anchored EGFP, fixed at indicated time points post biotin addition. The cells were fixed under non-permeabilising conditions and stained for surface GFP (yellow) with anti-GFP antibody. A delay in the trafficking of GPI-anchored EGFP (magenta) was observed, which was quantified by plotting the ratio of surface GFP to total GFP at different time points (N=3, n=70). Each point in the graph (**b**) represents mean ± SEM with the asterisks indicating a two-way ANOVA test significance value of p < 0.0001. Immunostaining NEU-treated cells for (**c**) E-cadherin (magenta), (**d**) β-catenin (yellow) and (**e**) α3-integrin showed an altered localisation of the cell-cell junction proteins. A pronounced distribution of the proteins in the cytoplasm (white arrow) and a diffused staining (green arrow) at the junctions was observed. The cytoplasmic distribution of the proteins was quantified by measuring the <u>C</u>orrected <u>T</u>otal <u>C</u>ell <u>F</u>luorescence (CTCF) of the cytoplasm. [CTCF = integrated density - (Area of selected cell x Mean fluorescence of background readings]. The CTCF for (f) E-cadherin (N=3, n=64), (g) β-catenin (N=3, n=40) and (h) α3-Integrin (N=3, n=56) has been represented as violin plots. Asterisks indicate Mann-Whitney U test significance values; **** p < 0.0001.

## Discussion

Previous studies from the lab have highlighted the role of DNA-PK in transforming breast epithelial cells (Anandi et al., 2017). Although these studies were done using 16-day breast acinar cultures, in this study, we observed changes in microtubule dynamics and trafficking as early as 48 hrs.

Changes in microtubule dynamics and dispersal of Golgi have been frequently associated with cancer progression (Bui et al., 2021; Petrosyan, 2015). Despite being thoroughly studied, the exact role of Golgi in cancer progression remains inconclusive. Golgi morphology is often perturbed in cancer cells that have resulted in an altered glycosylation signature and/or accelerated trafficking in cancer cells (Kellokumpu et al., 2002; Schultz et al., 2012; Wang et al., 2001). Cell migration, EMT, cell survival and proliferation, the nodal hallmarks of cancer, have been shown to regulate or be regulated by the Golgi apparatus (Bisel et al., 2008; Bui et al., 2021).

On the other hand, changes in microtubule dynamics have direct implications on cell migration and invasion (Etienne-Manneville, 2013; Lopes and Maiato, 2020; Wattanathamsan and Pongrakhananon, 2021). Microtubules in various cancer cell lines have been shown to have increased acetylation and detyrosination, which mark stable microtubules (Boggs et al., 2015; Mialhe et al., 2001). An increase in microtubule stabilising protein, Tau, has also been observed in metastatic breast cancer cell lines (Lei et al., 2020; Li et al., 2013). These observations have also been seen in breast tumour tissues from patients (Boggs et al., 2015; Whipple et al., 2010).

Invasion and migration require the cell to modulate the microtubule network and its interaction with the Golgi apparatus (Bui et al., 2021; Vinogradova et al., 2012). GDMTs have been linked to polarised trafficking to the cell’s leading edge, which is crucial for the initiation of migration and change in direction (Miller et al., 2009; Vinogradova et al., 2009). GDMTs are stable and marked by tubulin acetylation and are essential for maintaining Golgi structure (Vinogradova et al., 2012). Although evident when put together, no study has directly linked GDMTs to cancer progression. Our observations of microtubule stabilisation and GDMTs in MCF10CA1a show that microtubules are more stable and Golgi derived in malignant cell lines when compared to non-tumorigenic cells. Thus, this study is the first to provide an association between GDMTs, genomic instability and cancer progression.

Another aspect of microtubule regulation which is less understood is how the balance of centrosomal and non-centrosomal microtubules is maintained. Maintaining a balance between centrosomal and non-centrosomal microtubules decides the overall arrangement of the microtubule network. Altering this balance has often been seen as a mechanism to reorganise the microtubule network during cellular differentiation (Sanchez and Feldman, 2017). This reorganisation is mainly accomplished by inactivation or loss of centrosomal activity, which tips the balance to a more non-centrosomal array (Muroyama and Lechler, 2017; Nishita et al., 2017). A study by Rosa M Rios’s group in 2018 highlighted the plasticity of the interphase microtubule network and showed that centrosome dampens microtubule nucleation at Golgi and cytoplasm. In cells treated with centrinone to inhibit Plk4, the microtubule network was observed to be predominantly Golgi nucleated. It was also reported that cells with multiple centrosomes had little to no MT originating from Golgi (Gavilan et al., 2018). The authors suggest that centrosome-Golgi proximity might regulate the organisation of microtubules from the centrosome and Golgi. Golgi apparatus is associated with the centrosome, and the relevance of this proximity is unknown. A study in 2019 reported that Golgi-centrosome proximity was not essential for its function (Tormanen et al., 2019). Several studies deciphering the functional relevance of GDMTs have reported their importance for maintaining Golgi structure (Kodani and Sutterlin, 2009; Miller et al., 2009; Wu et al., 2016; Zhu and Kaverina, 2013). It is unknown whether the Golgi structure and its proximity to the centrosome regulate non-centrosomal microtubules that arise from Golgi. Our study provides a clue which hints that Golgi integrity might influence centrosomal control of GDMTs.

Based on findings from this and previous studies, we propose a model reporting a novel response to DNA damage **(Fig.7)**. DNA damage leads to the activation of DNA-PK, a DNA damage response protein. As previously reported, activation of DNA-PK leads to Golgi dispersal through the GOLPH3-MYO18A-F-actin axis (Dippold et al., 2009; Farber-Katz et al., 2014). Golgi dispersal leads to an increase in GDMTs through a mechanism yet to be elucidated. The microtubules nucleated at Golgi are stable, marked by enhanced tubulin acetylation, and resistant to nocodazole-induced depolymerisation. Changes in Golgi organisation and microtubule dynamics have been demonstrated to lead to defects in intracellular trafficking that may result in changes in cell polarity.

**Figure 7:**
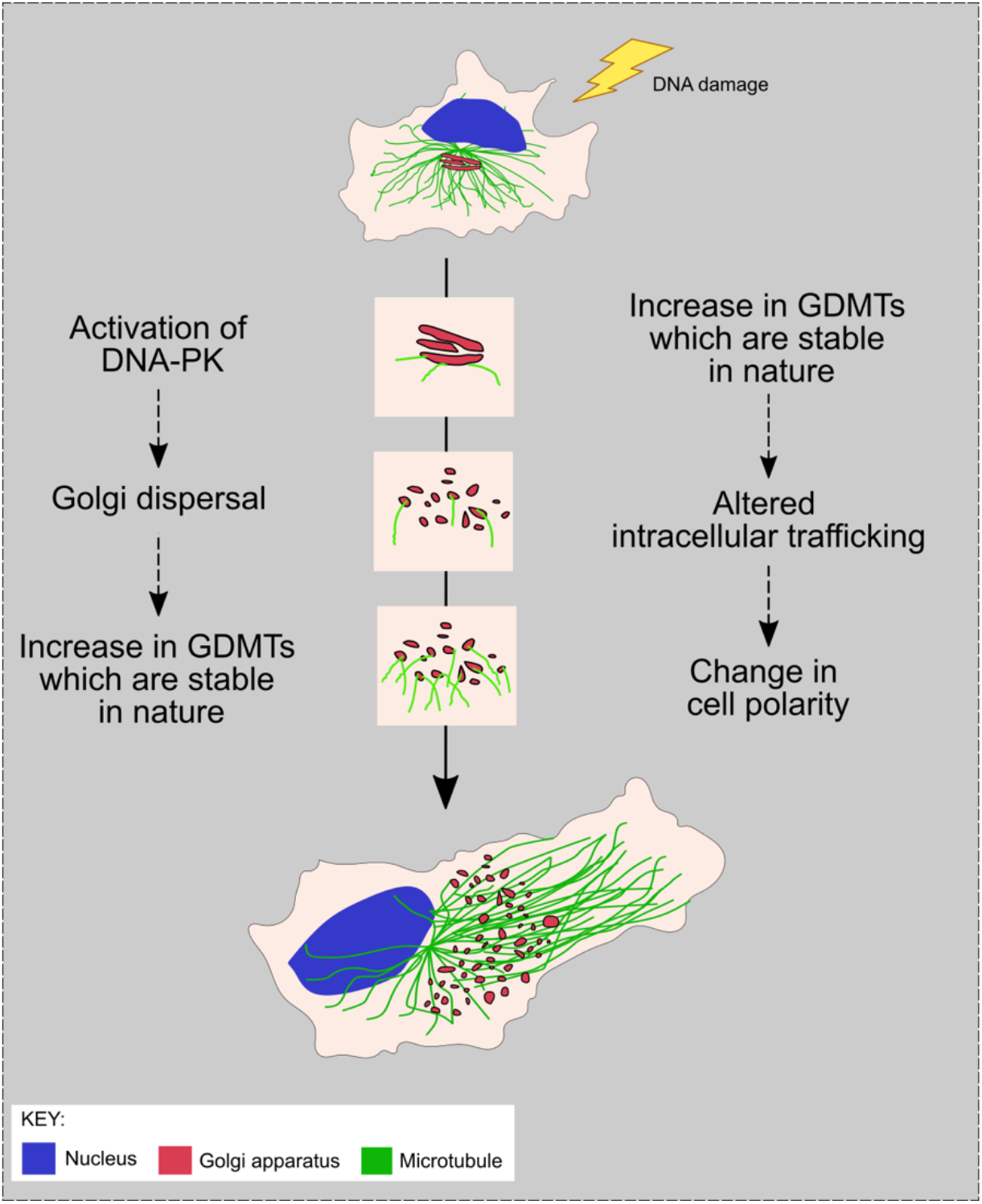
DNA damage leads to Golgi dispersal through the DNA-PK-GOLPH3-MYO18A axis. Golgi dispersal leads to an increase in GDMTs. Due to the difference in dynamics of GDMTs, the trafficking of proteins to the membrane gets altered, leading to mislocalisation of polarity proteins.

## Materials and Methods

### Cell line and culturing methods

MCF10A cells were a generous gift from Prof. Raymond C. Stevens (The Scripps Research Institute, La Jolla, CA). MCF10A and MCF10CA1a cell lines were cultured in DMEM containing high glucose without sodium pyruvate (Invitrogen) supplemented with 5% horse serum (Invitrogen), 100 units/ml penicillin/streptomycin (Invitrogen) and growth constituents as described in Anandi et al., 2017 (referred to as growth media in later text) (Anandi et al., 2017). HEK293 cells were grown in DMEM containing high glucose and sodium pyruvate (Lonza) supplemented with 10% foetal bovine serum (FBS) (Invitrogen) and 100 units/ml penicillin/streptomycin (Lonza) (referred to as complete DMEM in further text). Cells were cultured in tissue culture-treated 100mm or 60mm dishes (Eppendorf) in a 37°C humidified incubator with 5% CO_2_ (Eppendorf).

### Immunofluorescence

Cells were seeded at a density of 0.4 x10^5^ cells onto coverslips (Blue Star) in 24-well petri plates (Eppendorf). After appropriate drug treatment, the cells were fixed with 4% paraformaldehyde. The samples were washed with PBS-T (0.5% Triton-X100) and blocked with 10% FBS. Samples were then incubated in primary antibodies, diluted in 10% FBS, overnight at 4°C. Following incubation in primary antibody, cells were washed with PBS-T (0.5% Triton-X100) and incubated with secondary antibody (1:500) for 1 hour at room temperature. After a few washes with PBS-T (0.5% Triton-X100), the samples were incubated with Hoechst 33258 (Invitrogen) to stain the nucleus. Coverslips were mounted using 90% glycerol mounting media (with 20mM Tris, pH 8.0 and 0.5% propyl gallate). All images were obtained using Leica TCS SP8 X Confocal Microscope (Leica Microsystems). Post imaging, the maximum intensity projection of the stacks was used for analysis and has been represented in the figures.

### Microtubule regrowth assay

The cells were grown and treated with 2.5 μg/ml Nocodazole (Sigma) for 3 hrs at 37°C (Zhu and Kaverina, 2011). Following incubation, the cells were washed twice with ice-cold DPBS (Lonza) and supplemented with warm growth media. The microtubules were allowed to repolymerise at 37°C, fixed and stained for microtubule and Golgi and imaged on the Leica TCS SP8 X Confocal Microscope (Leica Microsystems).

### EB3 assay

MCF10A cells were seeded in 8-well chamber cover glass slides (Lab-Tek, Thermo Scientific) at a density of 0.6 x10^5^ cells and treated with 2 mM NEU (Sigma). After 24 hrs of treatment, the samples were transfected with the EB3-GFP construct. Transfections were performed using Lipofectamine 2000 (Invitrogen) in Opti-MEM (Invitrogen). The cells were then incubated for 4 hrs in the transfected medium, following which the media was changed to MCF10A growth media. The cells were imaged 24chrs after transfection in a live cell imaging chamber on the Leica TCS SP8 X Confocal Microscope (Leica Microsystems). Before imaging, the cells were shifted to L-15 medium (Lonza), and Hoechst 33342 was added to visualise the nucleus. Images were recorded at a frame rate of 1.5s per frame. The EB3 comets were manually tracked using the MTrackJ plugin (developed by E. Meijering, Biomedical Imaging Group, Erasmus Medical Center, Rotterdam) on Fiji/ImageJ (NIH)(Meijering et al., 2012).

### RUSH assay

To prepare the coverslips for low-adherent HEK293 cells, Matrigel (Sigma Aldrich) was diluted to 1:50 with DPBS (Lonza) and incubated for 4 hrs at 37°C. Subsequently, the cells were seeded at 0.6 x10^5^ cells and treated with 2 mM NEU. After 24 hrs of treatment, the cells were transfected with the RUSH constructs using bPEI25 (Sigma) using the protocol described in (Hsu and Uludag, 2012). DMEM (without serum) was used as the transfection medium instead of Opti-MEM to avoid the presence of biotin. The cells were then incubated for 4 hrs in the transfected medium, following which the media was changed to complete DMEM. 16 hrs posttransfection, RUSH assay was performed as described in (Boncompain et al., 2012; Boncompain and Perez, 2014). The cells were stained with Hoechst 33342 (Invitrogen) to mark the nucleus before mounting and visualised on the Leica TCS SP8 X Confocal Microscope (Leica Microsystems).

### Statistics

All graphs were plotted and analysed using Graph Pad Prism software (Graph Pad Software, La Jolla, CA, USA). All experimental groups were analysed by non-parametric methods except quantification of the RUSH assay. The significance of surface GFP/ total GFP in the RUSH assay was tested by two-way ANOVA. The Mann-Whitney test was used to analyse experimental groups with two samples. Groups with more than two samples were analysed using the Kruskal-Wallis test. Asterisks indicate significance values; **** p < 0.0001, ***p< 0.001, **p< 0.01, *p< 0.05 and ns – non-significant. The number of independent experiments performed and the number of cells analysed has been mentioned in the figure legends

## Acknowledgements

We would like to that Dr Aurnab Ghose (IISER Pune, India) for his valuable insights on the project and for also providing us with the EB3 construct and Latrunculin A. We thank Drs.Thomas Pucadyil and Richa Rikhy (IISER Pune, India) for their helpful suggestions. We also thank Dr Franck Perez (Institut Curie, Paris, France) for the RUSH plasmids and Dr Jomon Joseph (NCCS, India) for the K40 acetylated α-tubulin antibody. We thank Dr Libi Anandi and Snehal Bhatia from our laboratory for standardising some experiments. We also would like to acknowledge the IISER Pune Microscopy Facility for access to equipment and infrastructure.

## Author contributions

A.V. and M.L. conceived and conceptualised the project. A.V. designed, performed and analysed the experiments. A.V. and M.L. wrote the paper.

## References

Anandi, L., Chakravarty, V., Ashiq, K.A., Bodakuntla, S., and Lahiri, M. (2017). DNA-dependent protein kinase plays a central role in transformation of breast epithelial cells following alkylation damage. J Cell Sci 130, 3749–3763.

Bershadsky, A.D., and Gelfand, V.I. (1981). ATP-dependent regulation of cytoplasmic microtubule disassembly. Proceedings of the National Academy of Sciences of the United States of America 78, 3610–3613.

Bisel, B., Wang, Y., Wei, J.H., Xiang, Y., Tang, D., Miron-Mendoza, M., Yoshimura, S., Nakamura, N., and Seemann, J. (2008). ERK regulates Golgi and centrosome orientation towards the leading edge through GRASP65. The Journal of cell biology 182, 837–843.

Bodakuntla, S., Libi, A.V., Sural, S., Trivedi, P., and Lahiri, M. (2014). N-nitroso-N-ethylurea activates DNA damage surveillance pathways and induces transformation in mammalian cells. BMC cancer 14, 287.

Boggs, A.E., Vitolo, M.I., Whipple, R.A., Charpentier, M.S., Goloubeva, O.G., Ioffe, O.B., Tuttle, K.C., Slovic, J., Lu, Y., Mills, G.B., et al. (2015). alpha-Tubulin acetylation elevated in metastatic and basal-like breast cancer cells promotes microtentacle formation, adhesion, and invasive migration. Cancer Res 75, 203–215.

Boncompain, G., Divoux, S., Gareil, N., de Forges, H., Lescure, A., Latreche, L., Mercanti, V., Jollivet, F., Raposo, G., and Perez, F. (2012). Synchronization of secretory protein traffic in populations of cells. Nat Methods 9, 493–498.

Boncompain, G., and Perez, F. (2014). Synchronization of secretory cargos trafficking in populations of cells. Methods in molecular biology 1174, 211–223.

Bui, S., Mejia, I., Diaz, B., and Wang, Y. (2021). Adaptation of the Golgi Apparatus in Cancer Cell Invasion and Metastasis. Frontiers in cell and developmental biology 9, 806482.

Buschman, M.D., Rahajeng, J., and Field, S.J. (2015). GOLPH3 links the Golgi, DNA damage, and cancer. Cancer Res 75, 624–627.

Chabin-Brion, K., Marceiller, J., Perez, F., Settegrana, C., Drechou, A., Durand, G., and Pous, C. (2001). The Golgi complex is a microtubule-organizing organelle. Mol Biol Cell 12, 2047–2060.

Dippold, H.C., Ng, M.M., Farber-Katz, S.E., Lee, S.K., Kerr, M.L., Peterman, M.C., Sim, R., Wiharto, P.A., Galbraith, K.A., Madhavarapu, S., et al. (2009). GOLPH3 bridges phosphatidylinositol-4-phosphate and actomyosin to stretch and shape the Golgi to promote budding. Cell 139, 337–351.

Durant, S., and Karran, P. (2003). Vanillins--a novel family of DNA-PK inhibitors. Nucleic acids research 31, 5501–5512.

Etienne-Manneville, S. (2013). Microtubules in cell migration. Annu Rev Cell Dev Biol 29, 471–499.

Fang, Y.D., Xu, X., Dang, Y.M., Zhang, Y.M., Zhang, J.P., Hu, J.Y., Zhang, Q., Dai, X., Teng, M., Zhang, D.X., et al. (2011). MAP4 mechanism that stabilizes mitochondrial permeability transition in hypoxia: microtubule enhancement and DYNLT1 interaction with VDAC1. PloS one 6, e28052.

Farber-Katz, S.E., Dippold, H.C., Buschman, M.D., Peterman, M.C., Xing, M., Noakes, C.J., Tat, J., Ng, M.M., Rahajeng, J., Cowan, D.M., et al. (2014). DNA damage triggers Golgi dispersal via DNA-PK and GOLPH3. Cell 156, 413–427.

Fong, S., King, F., and Shtivelman, E. (2010). CC3/TIP30 affects DNA damage repair. BMC cell biology 11, 23.

Gavilan, M.P., Gandolfo, P., Balestra, F.R., Arias, F., Bornens, M., and Rios, R.M. (2018). The dual role of the centrosome in organizing the microtubule network in interphase. EMBO reports 19.

Geeraert, C., Ratier, A., Pfisterer, S.G., Perdiz, D., Cantaloube, I., Rouault, A., Pattingre, S., Proikas-Cezanne, T., Codogno, P., and Pous, C. (2010). Starvation-induced hyperacetylation of tubulin is required for the stimulation of autophagy by nutrient deprivation. The Journal of biological chemistry 285, 24184–24194.

Giannakakou, P., Sackett, D.L., Ward, Y., Webster, K.R., Blagosklonny, M.V., and Fojo, T. (2000). p53 is associated with cellular microtubules and is transported to the nucleus by dynein. Nat Cell Biol 2, 709–717.

Hsu, C.Y., and Uludag, H. (2012). A simple and rapid nonviral approach to efficiently transfect primary tissue-derived cells using polyethylenimine. Nature protocols 7, 935–945.

Hu, J.Y., Chu, Z.G., Han, J., Dang, Y.M., Yan, H., Zhang, Q., Liang, G.P., and Huang, Y.S. (2010). The p38/MAPK pathway regulates microtubule polymerization through phosphorylation of MAP4 and Op18 in hypoxic cells. Cellular and molecular life sciences: CMLS 67, 321–333.

Jackson, S.P., and Bartek, J. (2009). The DNA-damage response in human biology and disease. Nature 461, 1071–1078.

Kellokumpu, S., Sormunen, R., and Kellokumpu, I. (2002). Abnormal glycosylation and altered Golgi structure in colorectal cancer: dependence on intra-Golgi pH. FEBS letters 516, 217–224.

Kodani, A., and Sutterlin, C. (2009). A new function for an old organelle: microtubule nucleation at the Golgi apparatus. EMBO J 28, 995–996.

Lei, C., Yang, C., Xia, B., Ji, F., Zhang, Y., Gao, H., Xiong, Q., Lin, Y., Zhuang, X., Zhang, L., et al. (2020). Analysis of Tau Protein Expression in Predicting Pathological Complete Response to Neoadjuvant Chemotherapy in Different Molecular Subtypes of Breast Cancer. Journal of breast cancer 23, 47–58.

Li, Z.H., Xiong, Q.Y., Tu, J.H., Gong, Y., Qiu, W., Zhang, H.Q., Wei, W.S., Hou, Y.F., and Cui, W.Q. (2013). Tau proteins expressions in advanced breast cancer and its significance in taxane-containing neoadjuvant chemotherapy. Medical oncology 30, 591.

Lopes, D., and Maiato, H. (2020). The Tubulin Code in Mitosis and Cancer. Cells 9.

Ma, S., Rong, Z., Liu, C., Qin, X., Zhang, X., and Chen, Q. (2021). DNA damage promotes microtubule dynamics through a DNA-PK-AKT axis for enhanced repair. The Journal of cell biology 220.

Meijering, E., Dzyubachyk, O., and Smal, I. (2012). Methods for cell and particle tracking. Methods in enzymology 504, 183–200.

Mialhe, A., Lafanechere, L., Treilleux, I., Peloux, N., Dumontet, C., Bremond, A., Panh, M.H., Payan, R., Wehland, J., Margolis, R.L., et al. (2001). Tubulin detyrosination is a frequent occurrence in breast cancers of poor prognosis. Cancer Res 61, 5024–5027.

Miller, P.M., Folkmann, A.W., Maia, A.R., Efimova, N., Efimov, A., and Kaverina, I. (2009). Golgi-derived CLASP-dependent microtubules control Golgi organization and polarized trafficking in motile cells. Nat Cell Biol 11, 1069–1080.

Muroyama, A., and Lechler, T. (2017). Microtubule organization, dynamics and functions in differentiated cells. Development 144, 3012–3021.

Nakano, A., Kato, H., Watanabe, T., Min, K.D., Yamazaki, S., Asano, Y., Seguchi, O., Higo, S., Shintani, Y., Asanuma, H., et al. (2010). AMPK controls the speed of microtubule polymerization and directional cell migration through CLIP-170 phosphorylation. Nat Cell Biol 12, 583–590.

Nishita, M., Satake, T., Minami, Y., and Suzuki, A. (2017). Regulatory mechanisms and cellular functions of non-centrosomal microtubules. Journal of biochemistry 162, 1–10.

Parker, A.L., Kavallaris, M., and McCarroll, J.A. (2014). Microtubules and their role in cellular stress in cancer. Frontiers in oncology 4, 153.

Petrosyan, A. (2015). Onco-Golgi: Is Fragmentation a Gate to Cancer Progression? Biochemistry & molecular biology journal 1.

Poruchynsky, M.S., Komlodi-Pasztor, E., Trostel, S., Wilkerson, J., Regairaz, M., Pommier, Y., Zhang, X., Kumar Maity, T., Robey, R., Burotto, M., et al. (2015). Microtubule-targeting agents augment the toxicity of DNA-damaging agents by disrupting intracellular trafficking of DNA repair proteins. Proc Natl Acad Sci U S A 112, 1571–1576.

Ryu, N.M., and Kim, J.M. (2020). The role of the alpha-tubulin acetyltransferase alphaTAT1 in the DNA damage response. J Cell Sci 133.

Sanchez, A.D., and Feldman, J.L. (2017). Microtubule-organizing centers: from the centrosome to non-centrosomal sites. Curr Opin Cell Biol 44, 93–101.

Schultz, M.J., Swindall, A.F., and Bellis, S.L. (2012). Regulation of the metastatic cell phenotype by sialylated glycans. Cancer metastasis reviews 31, 501–518.

Skoufias, D.A., Burgess, T.L., and Wilson, L. (1990). Spatial and temporal colocalization of the Golgi apparatus and microtubules rich in detyrosinated tubulin. J Cell Biol 111, 1929–1937.

Thyberg, J., and Moskalewski, S. (1993). Relationship between the Golgi complex and microtubules enriched in detyrosinated or acetylated alpha-tubulin: studies on cells recovering from nocodazole and cells in the terminal phase of cytokinesis. Cell and tissue research 273, 457–466.

Tormanen, K., Ton, C., Waring, B.M., Wang, K., and Sutterlin, C. (2019). Function of Golgi-centrosome proximity in RPE-1 cells. PloS one 14, e0215215.

Vinogradova, T., Miller, P.M., and Kaverina, I. (2009). Microtubule network asymmetry in motile cells: role of Golgi-derived array. Cell Cycle 8, 2168–2174.

Vinogradova, T., Paul, R., Grimaldi, A.D., Loncarek, J., Miller, P.M., Yampolsky, D., Magidson, V., Khodjakov, A., Mogilner, A., and Kaverina, I. (2012). Concerted effort of centrosomal and Golgi-derived microtubules is required for proper Golgi complex assembly but not for maintenance. Molecular biology of the cell 23, 820–833.

Wang, F., Goto, M., Kim, Y.S., Higashi, M., Imai, K., Sato, E., and Yonezawa, S. (2001). Altered GalNAc-alpha-2,6-sialylation compartments for mucin-associated sialyl-Tn antigen in colorectal adenoma and adenocarcinoma. The journal of histochemistry and cytochemistry: official journal of the Histochemistry Society 49, 1581–1592.

Wattanathamsan, O., and Pongrakhananon, V. (2021). Post-translational modifications of tubulin: their role in cancers and the regulation of signaling molecules. Cancer gene therapy.

Wehland, J., Henkart, M., Klausner, R., and Sandoval, I.V. (1983). Role of microtubules in the distribution of the Golgi apparatus: effect of taxol and microinjected anti-alpha-tubulin antibodies. Proc Natl Acad Sci U S A 80, 4286–4290.

Whipple, R.A., Matrone, M.A., Cho, E.H., Balzer, E.M., Vitolo, M.I., Yoon, J.R., Ioffe, O.B., Tuttle, K.C., Yang, J., and Martin, S.S. (2010). Epithelial-to-mesenchymal transition promotes tubulin detyrosination and microtentacles that enhance endothelial engagement. Cancer research 70, 8127–8137.

Williams, T., Courchet, J., Viollet, B., Brenman, J.E., and Polleux, F. (2011). AMP-activated protein kinase (AMPK) activity is not required for neuronal development but regulates axogenesis during metabolic stress. Proceedings of the National Academy of Sciences of the United States of America 108, 5849–5854.

Wu, J., de Heus, C., Liu, Q., Bouchet, B.P., Noordstra, I., Jiang, K., Hua, S., Martin, M., Yang, C., Grigoriev, I., et al. (2016). Molecular Pathway of Microtubule Organization at the Golgi Apparatus. Dev Cell 39, 44–60.

Xu, Z., Schaedel, L., Portran, D., Aguilar, A., Gaillard, J., Marinkovich, M.P., Thery, M., and Nachury, M.V. (2017). Microtubules acquire resistance from mechanical breakage through intralumenal acetylation. Science 356, 328–332.

Yoon, S.O., Shin, S., and Mercurio, A.M. (2005). Hypoxia stimulates carcinoma invasion by stabilizing microtubules and promoting the Rab11 trafficking of the alpha6beta4 integrin. Cancer research 65, 2761–2769.

Zhu, X., and Kaverina, I. (2011). Quantification of asymmetric microtubule nucleation at subcellular structures. Methods in molecular biology 777, 235–244.

Zhu, X., and Kaverina, I. (2013). Golgi as an MTOC: making microtubules for its own good. Histochemistry and cell biology 140, 361–367.

