## Supplementary Information for "DNA damage leads to microtubule stabilisation through an increase in Golgi-derived microtubules"

**Table 1: Reagents table**

| Reagent | Source | Dilution/Concentration |
|---|---|---|
| Antibodies and dyes | | |
| α-Tubulin | Sigma | 1:500 |
| α-Tubulin | Abcam | 1:500 |
| GM130-FITC conjugated | BD Biosciences | 1:100 |
| GM130 | Abcam | 1:100 |
| Acetylated a-Tubulin (K40) | Sigma | 1:500 |
| pDNA-PK (Thr 2609) | Abcam | 1:200 |
| Anti-GFP antibody | Abcam | 1:1000 |
| Hoechst 33258 | Invitrogen | 1:10,000 |
| Hoechst 33342 | Invitrogen | 1:1000 |
| Phalloidin (Alexa Fluor 568 or 633 conjugated) | Invitrogen | 1:100 |
| Alexa Fluor 488, 568 or 633 conjugated secondary antibodies | Invitrogen | 1:500 |
| Chemicals | | |
| N-nitroso-N-ethylurea (NEU) | Sigma | 2 mM (in MCF10A) and 0.5 mM (in HEK293) |
| DMNB | Tocris | 25mM |
| Latrunculin A | Sigma | 250nM |
| Biotin | Sigma | 40mM |

**SUPPLEMENTARY FIGURES**

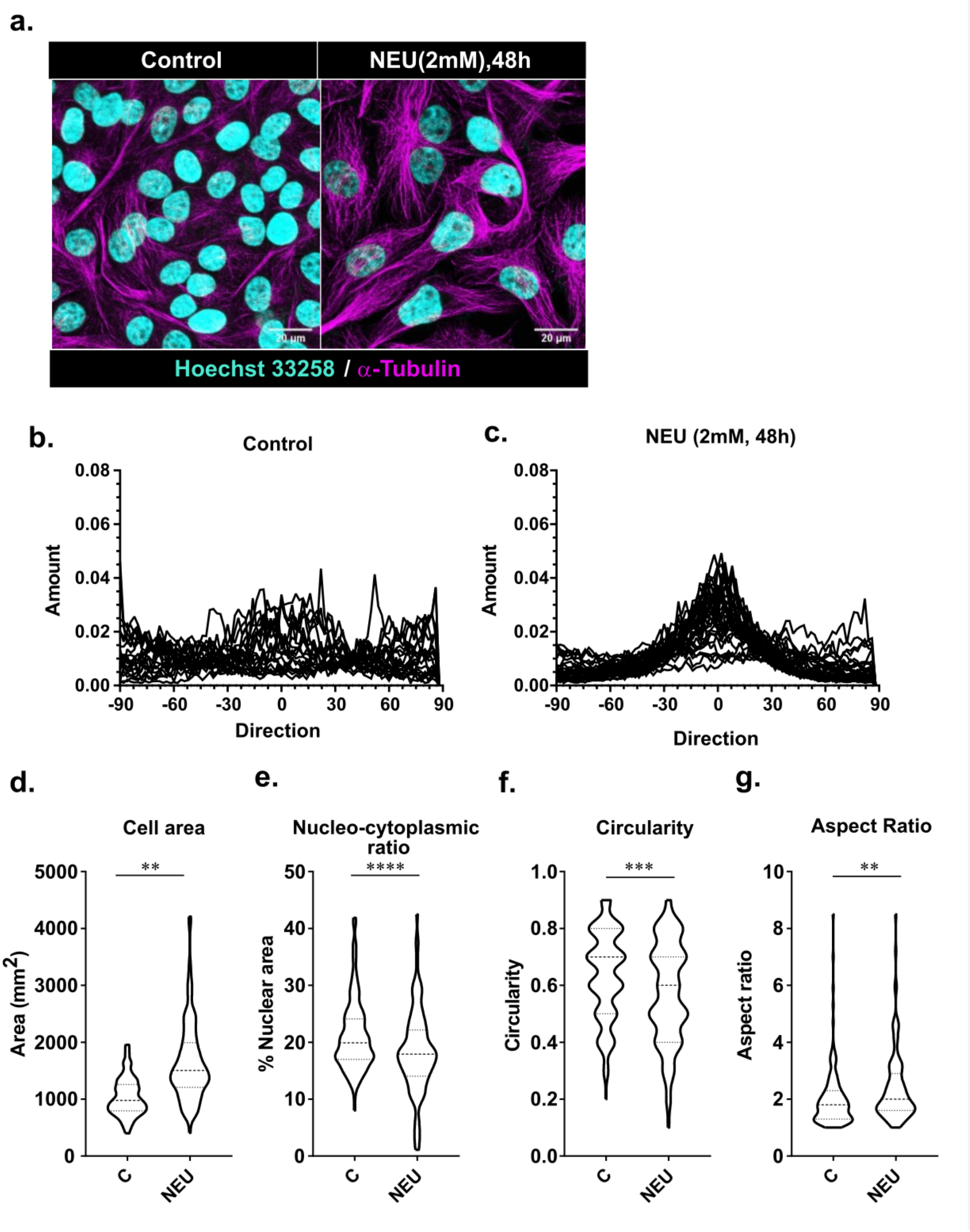


**Supp Fig 1: NEU treatment to MCF10A cells led to changes in cell morphology (a)** MCF10A cells treated with NEU (2mM, 48h) showed a parallel arrangement of microtubules (magenta) that was quantified in (**b and c)** to depict microtubule orientation in cells treated with and without NEU (N=3, n=30). ImageJ directionality tool was used to generate the orientation plots. NEU-induced DNA damage led to a significant increase in **(d)** cell area, **(e)** nucleo-cytoplasmic ratio, (**f**) circularity and (**g**) aspect ratio, which were measured using ImageJ (N=3, n=160). Asterisks indicate Mann-Whitney U test significance values; **** p < 0.0001; *** p< 0.001; ** p< 0.01.

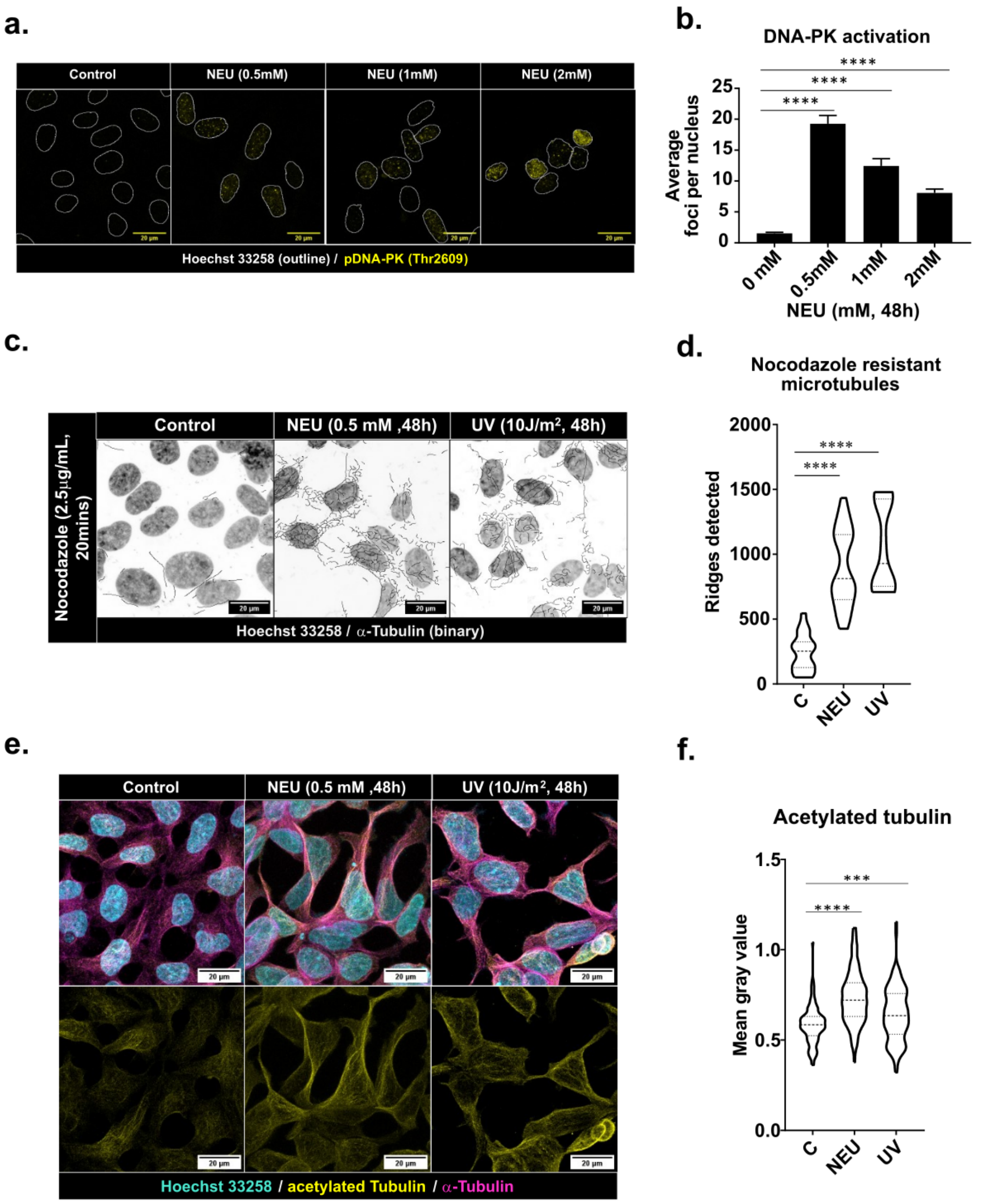


**Supp Fig 2: HEK293 cells showed an increase in microtubule stability post DNA damage**. HEK293 cells were treated with different doses of NEU and stained for pDNA-PK (T2609). (**a)** Activation of DNA-PK was observed in NEU-treated cells by the presence of foci (yellow) in the nucleus stained with Hoechst 33258 (white outline). (**b)** Quantification of the number of pDNA-PK foci per nucleus showed significant increases in DNA-PK activation upon NEU treatment (N=3, n=200). (**c**)Treatment with nocodazole revealed a subset of microtubules in cells treated with DNA damage resistant to depolymerisation. Shown are the binary images of the subset of microtubules and the nucleus. (**d**) Quantification of the microtubule subset using ImageJ ridge detection tool (N=3, n>10 fields per treatment). (**e**) HEK293 cells treated with NEU (0.5 mM, 48 hrs) and UV (10 J/m$^2$) stained for α-tubulin (magenta), acetylated-tubulin (yellow), and Hoechst 33258 (cyan) showed an increase in tubulin acetylation post DNA damage. (**f**) Quantification of the acetylated tubulin represented as violin plots (N=3, n=135). Asterisks indicate Mann-Whitney U test significance values; **** $p < 0.0001$; *** $p < 0.001$.

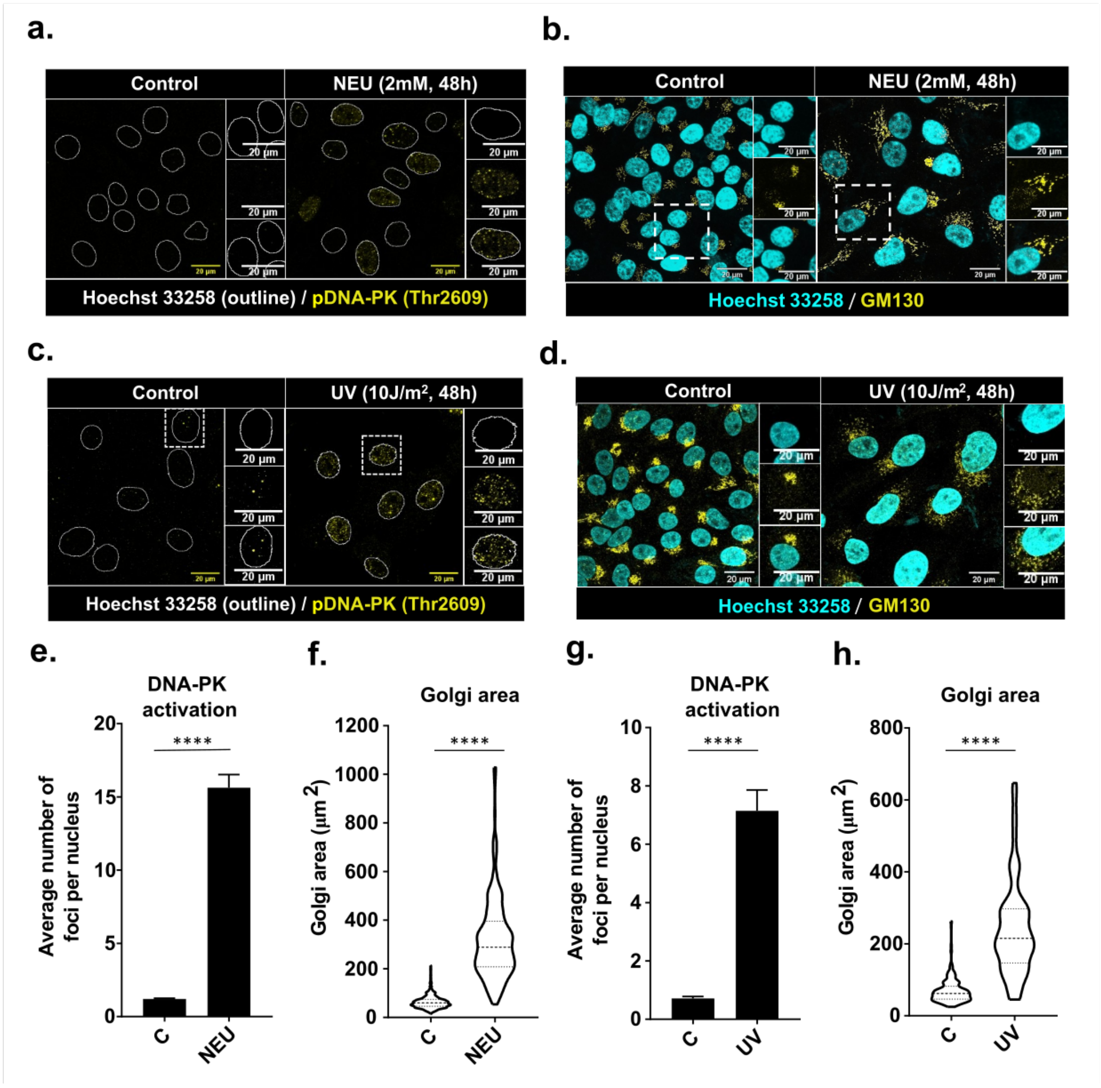


**Supp Fig 3: MCF10A cells treated with NEU (2mM) and UV (10J/m$^2$) showed activation of DNA-PK and Golgi dispersal.** (**a** and **c**) Cells stained for pDNA-PK (yellow) and Hoechst 33258 nuclear outline (white), following 2 mM NEU for 48 hrs (N=3, n=200) and 10 J/m$^2$ UV for 48 hrs (N=3, n=180) treatments, respectively. (**b** and **d**) Cells stained with GM130, a cis-Golgi marker (yellow) and Hoechst 33258 (cyan) for the nucleus, following NEU and UV treatments, respectively. (**e** and **g**) Quantifying the number of pDNA-PK foci observed in the nucleus of the cells following DNA damage treatment. (**f** and **h**) Violin plots representing the Golgi area of cells treated with NEU and UV. Asterisks indicate Mann-Whitney U test significance values; **** $p < 0.0001$ (N=3, n=210 cells).

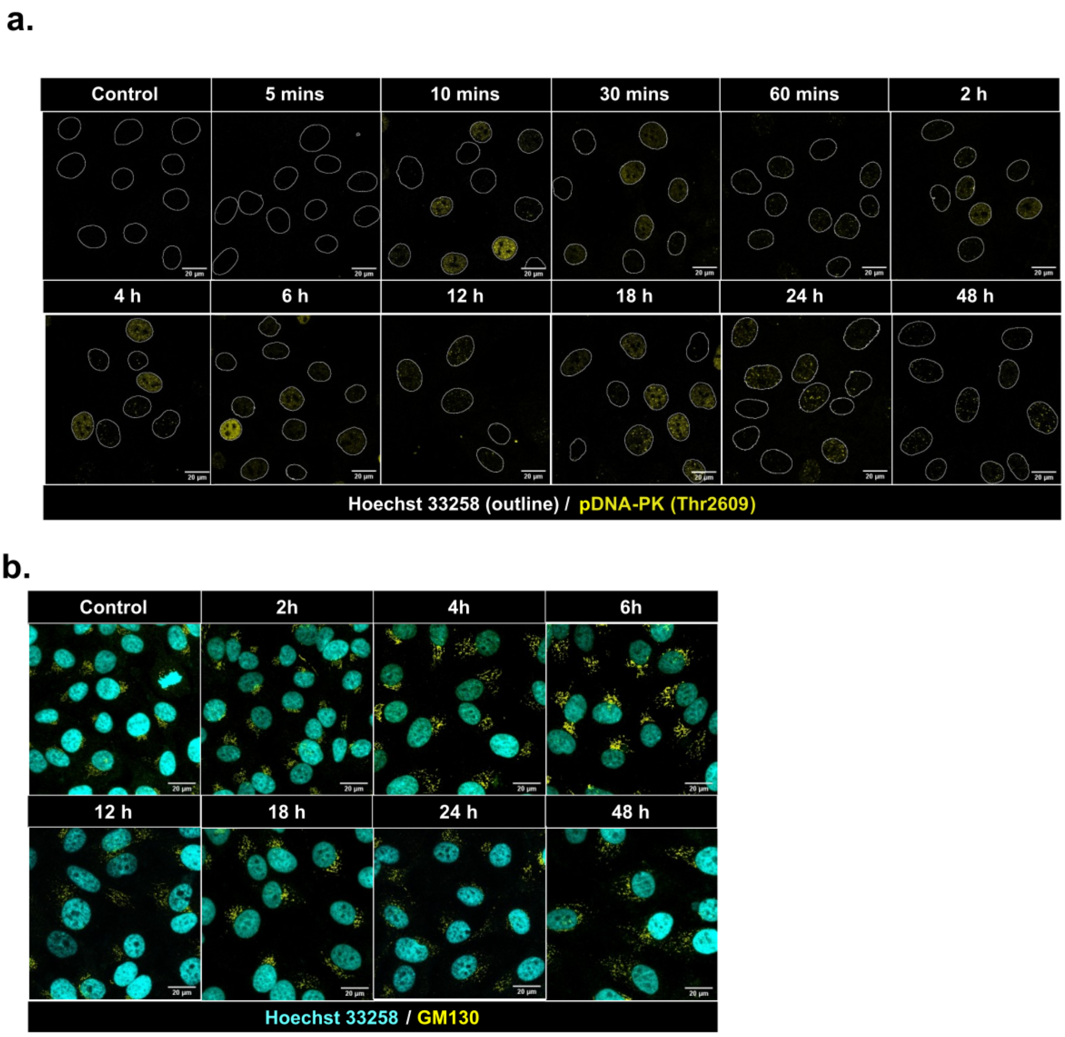


**Supp Figure 4: Activation of DNA-PK precedes Golgi dispersal following NEU treatment.** Cells were treated with 2 mM NEU for different durations, and (**a**) DNA-PK activation was observed from 10 mins, which remained active till 48 hrs. while (**b**) Golgi started to disperse from 4hrs onwards.

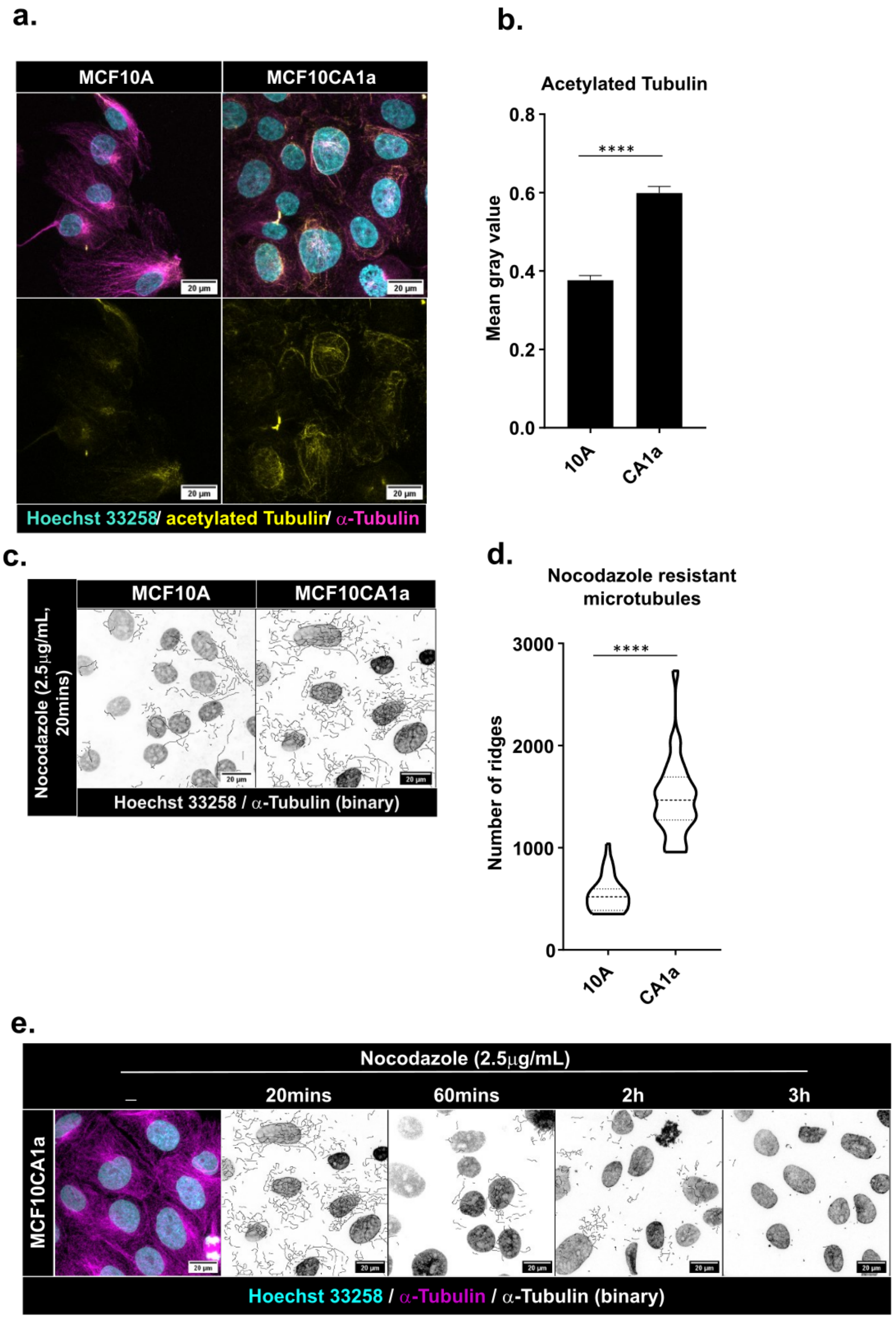


**Supp Fig 5: Microtubule stabilisation in MCF10CA1a.** MCF10A and MCF10CA1a cells were stained with α-tubulin (magenta), acetylated tubulin (yellow) and Hoechst 33258 (cyan). Increased tubulin acetylation was observed in MCF10CA1a compared to MCF10A, which was quantified and plotted in (**b.**) (N=3, n=150 cells). An increase in nocodazole-resistant microtubules was also observed in MCF10CA1a cells. Shown in (**c.**) are binary images of microtubules that fail to depolymerise post nocodazole treatment, which were quantified using the ridge detection tool and plotted in (**d.**) (N=3, n>10 fields per sample). Asterisks indicate Mann-Whitney U test significance values; **** $p < 0.0001$. **e.** A nocodazole dose of 2.5µg/mL for 3h led to the complete depolymerisation of microtubules in MCF10CA1a cells. Representative images of MCF10CA1a cells incubated with 2.5µg/mL of nocodazole for 20mins, 60mins, 2h and 3h is shown.

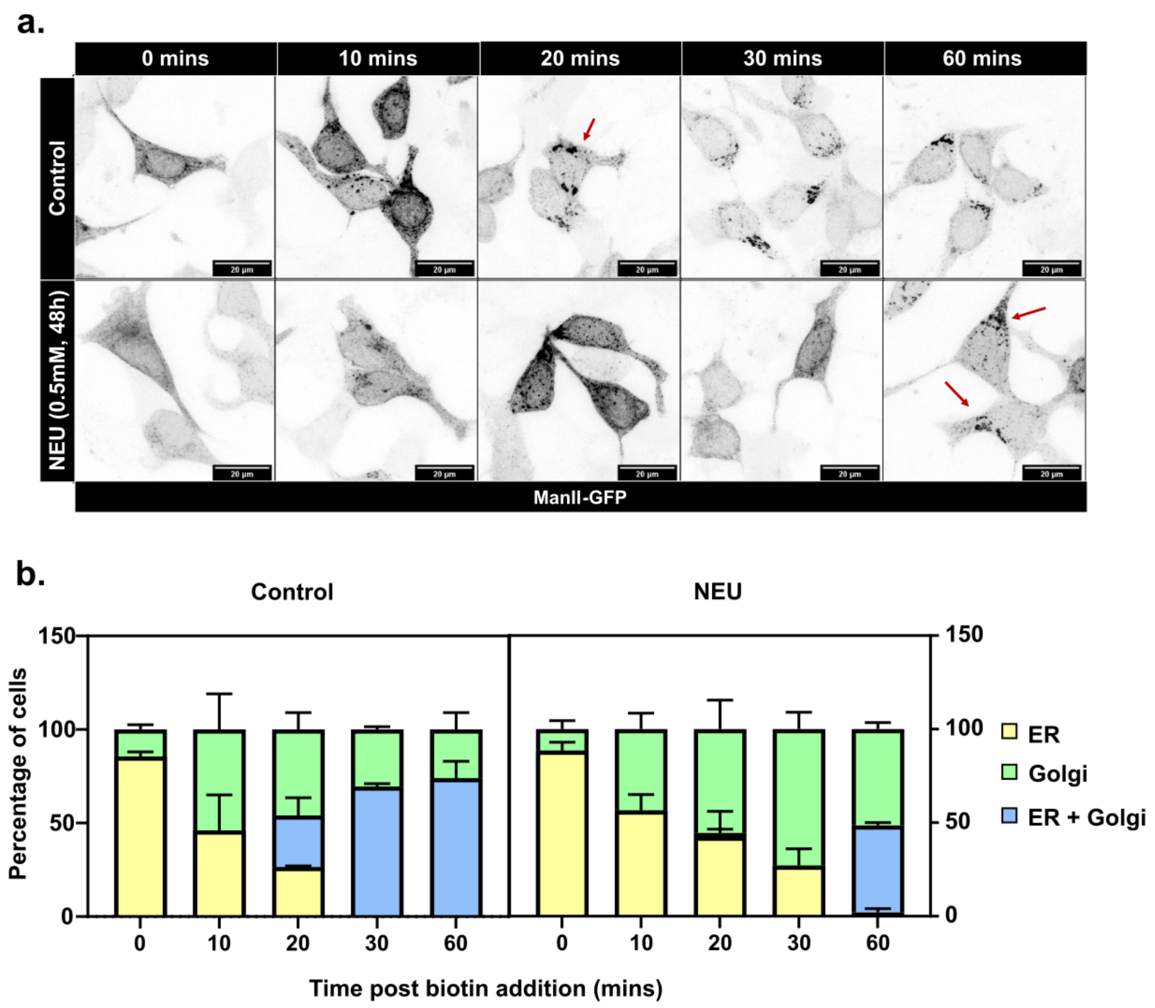


**Supp Fig 6: DNA damage led to altered intracellular trafficking.** (**a**) Delayed trafficking of ManII-GFP to Golgi apparatus was observed upon induction of DNA damage in HEK 293 cells treated with 0.5 mM NEU for 48 hrs. Arrows depict the cargo that had reached the Golgi apparatus. (**b**) The graph represents the percentage of cells showing GFP signal in ER (yellow), Golgi (blue) or both (green) at the indicated time points post biotin addition (N=3, n=60).
